# Genome-wide association study identifies 17 new loci influencing concentrations of circulating cytokines and growth factors

**DOI:** 10.1101/045005

**Authors:** AV Ahola-Olli, P Würtz, AS Havulinna, K Aalto, N Pitkänen, T Lehtimäki, M Kähönen, LP Lyytikäinen, E Raitoharju, I Seppälä, AP Sarin, S Ripatti, A Palotie, M Perola, JS Viikari, S Jalkanen, M Maksimow, V Salomaa, M Salmi, J Kettunen, OT Raitakari

**Affiliations:** Research Centre of Applied and Preventive Cardiovascular Medicine, University of Turku, Turku, Finland.; Computational Medicine, Center for Life Course Health Research, University of Oulu, Oulu, Finland; National Institute of Health and Welfare, Helsinki, Finland; Medicity Research Laboratory, Department of Medical Microbiology and Immunology, University of Turku, Turku, Finland; The Department of Clinical Chemistry, Fimlab Laboratories, Pirkanmaa Hospital District, School of Medicine, University of Tampere, Tampere, Finland; The Department of Clinical Physiology, Tampere University Hospital, Tampere, Finland; Institute for Molecular Medicine Finland FIMM, University of Helsinki, Helsinki, Finland; National Institute for Health and Welfare, Helsinki, Finland; Institute for Molecular Medicine Finland FIMM, University of Helsinki, Helsinki, Finland; National Institute for Health and Welfare, Helsinki, Finland; The Department of Public Health, University of Helsinki, Helsinki, Finland; Analytic and Translational Genetics Unit, Department of Medicine, Massachusetts General Hospital, Boston, MA; Program in Medical and Population Genetics, Broad Institute of MIT and Harvard, Cambridge, MA; Stanley Center for Psychiatric Research, Broad Institute of MIT and Harvard, Cambridge, MA; Institute for Molecular Medicine Finland, University of Helsinki, Helsinki, Finland; Psychiatric and Neurodevelopmental Genetics Unit, Department of Psychiatry, Massachusetts General Hospital, Boston, MA; Department of Neurology, Massachusetts General Hospital, Boston, MA; The Department of Medicine, University of Turku and Division of Medicine, Turku University Hospital, Turku, Finland; NMR Metabolomics Laboratory, School of Pharmacy, University of Eastern Finland, Kuopio, Finland; Biocenter Oulu, University of Oulu, Finland; The Department of Clinical Physiology and Nuclear Medicine, Turku University Hospital, Turku, Finland

## Abstract

Circulating cytokines and growth factors are regulators of inflammation and have been implicated in autoimmune and metabolic diseases. In this genome-wide association study (GWAS) up to n=8,293 Finns we identified 27 loci with genome-wide association (P-value<1.2×10^-9^) for one or more cytokines, including 17 unidentified in previous GWASes. Fifteen of the associated SNPs had expression quantitative trait loci in whole blood. We provide strong genetic instruments to clarify the causal roles of cytokine signaling and upstream inflammation in immune-related and other chronic diseases. We further link known autoimmune disease variants including Crohn's disease, multiple sclerosis and ulcerative colitis with new inflammatory markers, which elucidate the molecular mechanisms underpinning these diseases and suggest potential drug targets.

---

Genome-wide association studies (GWAS) have identified multiple loci for inflammatory diseases^1,2^. However, the biochemical pathways underlying the link from locus to complex disease have often remained elusive. Cytokines and growth factors (hereafter cytokines) are inflammatory regulators, and therefore important intermediate phenotypes for inflammatory diseases.^3^ These intermediate phenotypes can be exploited in GWASes to elucidate the biochemical pathways underlying the link from locus to disease susceptibility. Cytokines have been implicated in the progression of inflammatory bowel diseases, multiple sclerosis, atherosclerosis and in various forms of cancer. Here, we studied the genetic basis for blood levels of multiple cytokines to gain insights to the molecular intermediates and causal pathways related to inflammatory diseases.

The genetic basis of inflammatory diseases can inform drug development to prioritise new pharmaceutical targets. For example, an individual who has inherited a cytokine signaling disrupting allele is analogous to a participant in randomized trial receiving active drug designed to inhibit signaling via the pertinent cytokine. Cytokines are attractive targets since they exert their effects via cell surface receptors that are readily druggable. Characterization of the genetic underpinnings of cytokine signaling pathways is important for drug development, because pharmaceutical targets with human genetic support have twice the probability for eventual regulatory approval compared to non-supported targets^4^. Furthermore, since inflammatory diseases often share common pathology, the indications of approved drugs can often be expanded^5^. For example, a tumor necrosis factor alpha (TNF-α) antibody adalimumab has FDA approval for rheumatoid arthritis, juvenile idiopathic arthritis, ankylosing spondylitis, psoriatic arthritis and Crohn’s disease, although initially studied for rheumatoid arthritis^6–8^. Therefore, to gain insights into the molecular intermediates and causal pathways related to autoimmune and metabolic diseases, we studied the genetic basis for circulating levels of 41 cytokines. In addition to informing drug development, our results provide novel information on the genetic regulation of normal physiological variation of cytokines among healthy individuals.

## Results

We performed GWAS for the circulating concentrations of 41 cytokines; listed in Supplementary **Table 1**. The GWAS meta-analysis included up to 8,293 Finnish individuals from three independent population cohorts: the Cardiovascular Risk in Young Finns Study (YFS), FINRISK1997 and FINRISK2002. Study cohort characteristics are reported in **Supplementary Table 1**. On average, the YFS participants are younger than the FINRISK1997 and FINRISK2002 participants (37 vs. 60 years). Correlations between the cytokine levels are shown in **Supplementary Figure 1**. Four cytokines (VEGF, IL12p70, IL13 and IL10) form a tightly correlated cluster. Most of the cytokines included in the study are positively correlated with each other.

**Table 1.**
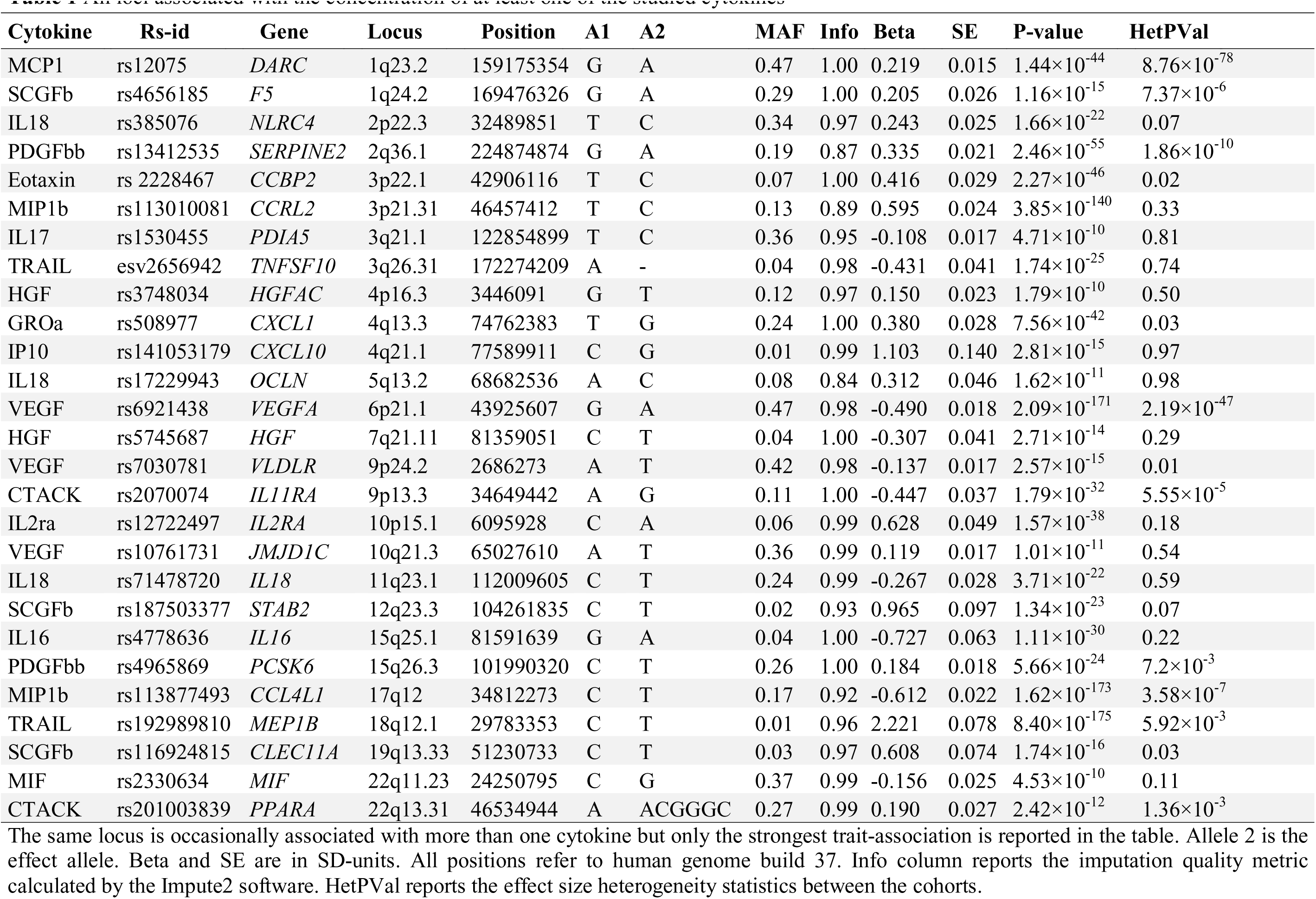
All loci associated with the concentration of at least one of the studied cytokines

### GWAS identifies 27 loci associated with circulating cytokines

Using an additive genetic model adjusted for the ten first genetic principal components, age, sex and body mass index, we tested for univariate associations between 10.7 million genetic polymorphisms and 41 cytokine concentrations. Genotype imputation was performed based on reference haplotypes provided by the 1000 Genomes Project September 2013 release^9^. The meta-analyses identified 27 genome-wide significant loci (*P*<1.2×10^-9^ accounting for 41 measures); 17 of which have not been associated with cytokines, C-reactive protein or white blood cell count in previous GWASes (**Table 1 and Figure 1**). With the aforementioned threshold for genome-wide significance and sample size, we had 84 % power to detect an association with effect size of 0.11 standard deviations per one additional copy of effect allele assuming minor allele frequency (MAF) of 50%. The potential candidate genes of the 27 genetic variants are also reported in **Table 1**. Results from the random effect meta-analysis for heterogenic single nucleotide polymorphisms (SNPs) (HetPval < 0.1) are reported in **Supplementary Table 2**. Effect estimates from the random effect models were coherent with effect estimates from the fixed effect models; however, for ten of the heterogenic loci the P-value did not reach the genome-wide significance (1.2x10^-9^) with the random effect model. Complete summary statistics from meta-analyses are available at (http://www.will_be_announced_later.com). Cytokine concentration means per allele in absolute concentration units for lead SNPs are reported in **Supplementary Table 3** and genomic lambda values in **Supplementary Table 4**. The lambda values ranged from 0.98 to 1.02 indicating that there was no overall inflation in the test statistics.

**Figure 1.**
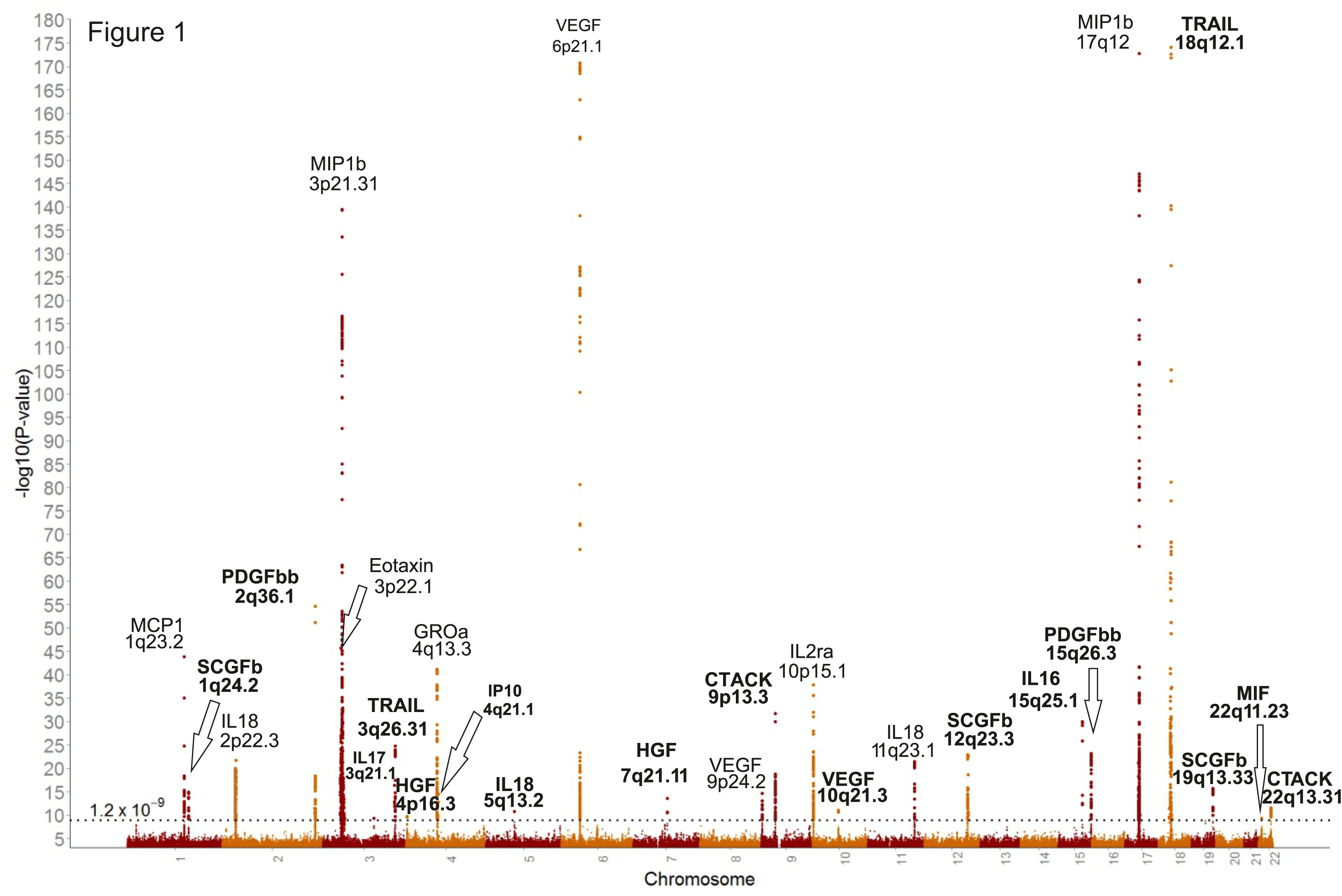
Combined Manhattan plot of meta-analysis results from the GWAS of 41 circulating cytokines. Loci not previously associated with cytokine concentration in GWAS are bolded. The horizontal dashed line indicates the genome-wide significance threshold (*P*<1.2×10^-9^) accounting for the number of cytokines tested in the study.

Novel loci with large effects included rs192989810 from 18q12.1 for TNF-related apoptosis inducing ligand (TRAIL) with β=2.2 SD at MAF of 1%. This SNP is located within *MEP1B*, and leukocytes from *MEP1B-/-* mice have been shown to have impaired migration in response to monocyte chemotactic protein-1 (MCP1) and macrophage inflammatory protein-1α (MIP1a)^10^. Additional low-frequency variants with large effects included rs141053179 (β=1.1 SD) for interferon gamma-induced protein 10 (IP10), rs116924815 (β=0.61 SD) for stem cell growth factor beta (SCGFb), and rs4778636 (β=-0.73 SD) for interleukin-16 (IL16). Novel common variants (MAF>5%) with large effects (β up to ‐0.49) included rs2070074 and rs13412535 for Cutaneous T-cell attracting (CTACK) and platelet derived growth factor BB (PDGFbb). In addition to discovering new loci, our results confirmed associations for 10 genome-wide significant loci previously identified. These include two loci with interleukin-18 (IL18) concentration (2p22.3 and 11q23.1)^11,12^. We also replicated two loci previously associated with vascular endothelial growth factor (VEGF; 6p21.1 and 9p24.2)^13^ and one additional locus previously associated with monocyte chemotactic protein-1 (MCP1; 1q23.2)^14^. Manhattan plots, QQ-plots and detailed plots from heterogeneity for each cytokine are provided in the **Supplementary note**. For 6p21.1 associated with vascular endothelial growth factor (VEGF), the association signal was narrow and hereby suggested only few causal SNP candidates (Supplementary note, page 55). After conditioning on rs6921438, the subsequent lead SNP was rs12214617 which is located 46kB from rs6921438. According to Ensembl, the secondary association signal rs12214617 is located on the promoter flanking region in various cell types including monocytes and HepG2 cells, thus suggesting a potential role in gene regulation. A previous GWAS of VEGF levels identified the same lead SNP associated with VEGF concentration in 6p21.1^13^. Our results confirm this association but the secondary associations signal was different from ours, potentially due to different imputation reference. For some of the genetic loci associated with cytokines the association signal is much wider, which makes the identification of the potential functional variant more challenging. For instance, the signal spans approximately 600kB for IL18-associated 2p22.3. The association peak for PDGFbb in 2q36.1 is mainly formed by only two variants (rs13412535 and rs68066031) in strong linkage disequilibrium. However, after conditioning on rs13412535 the peak is lost. According to Ensembl, the rs13412535 is located in a transcription factor binding cite and DNase peak indicating accessible chromatin. Thus, rs13412535 is likely affecting to transcription factor binding and gene expression of *SERPINE2*.

Four of the identified loci were associated with the concentration of more than one cytokine (1q23.2, 6p21.1, 3p22.1, and 10q21.3). The 1q23.2 harbors rs12075, which is genome-wide significant for three cytokines: eotaxin, MCP1 and growth-regulated oncogene-α (GROa). The lead SNP is located within *DARC*, which encodes a receptor for multiple cytokines and is also a human erythrocyte blood group antigen^15,16^. This suggests that the effect of rs12075 on the three cytokines is not directly mediated via any of the three pertinent cytokines, but rather through altering binding of these cytokines to the DARC receptor (**Fig. 2a**). Similarly, the effect of rs2228467 at 3p22.1 on eotaxin and MCP1 appears to be mediated via chemokine-binding protein 2 (*CCBP2*), rather than directly via eotaxin or MCP1 (**Fig. 2b**). In contrast, a variant at 6p21.1 (rs6921438) is associated with concentration of five cytokines (VEGF, IL12p70, IL7, IL10, and IL13). This locus contains a *VEGFA* gene encoding VEGF, which suggest that VEGF might regulate the concentration of other cytokines associated with rs6921438. To test this directional mediation, we performed Mendelian randomization analysis using a genetic score of rs6921438 and rs12214617 (variants at 6p21.1 independently associated with VEGF) as the instrumental variable^17^. The observational and causal effect estimates were matching each other, supporting the role of VEGF as an upstream regulator of IL12p70, IL7, IL10, and IL13 (**Fig. 2c-d**). Variants in 10q21.3 were associated with VEGF and IL12p70. The lead SNP within this locus is rs10761731, which has the smallest association test P-value for VEGF. The ratio of the IL12p70 effect size divided by the effect size of VEGF remains approximately constant in both loci (6p21.1 and 10q21.3) suggesting tight regulatory effect of VEGF on IL12p70 concentration.

**Figure 2.**
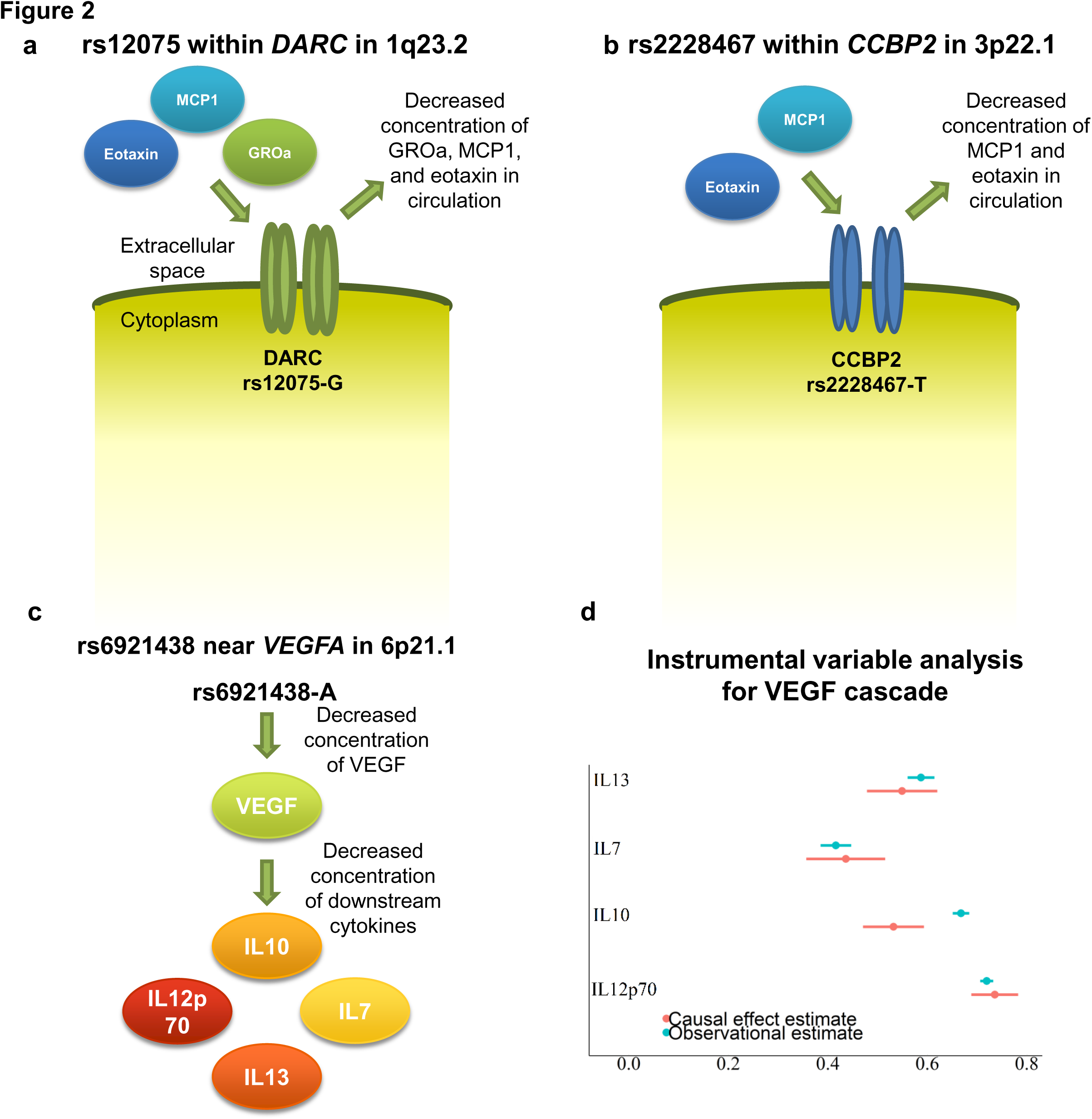
SNPs genome-widely associated with more than one cytokine. Variant rs12075-A is associated with increased concentrations of eotaxin, monocyte chemotactic protein-1 (MCP1) and growth regulated oncogene-a (GROa). In addition, rs12075-A is associated with increased *DARC* mRNA. Since DARC is a cell surface receptor, the cluster effect on the three cytokines is unlikely to be mediated (in a cascade) through any of the three cytokines associated with rs12075 (Panel a). Similarly, effect of rs2228467 is mediated through a cell surface receptor (*CCBP2*) (Panel b). Variant rs6921438 in 6p21.1 is located near *VEGFA*. This suggests that VEGF is causally regulating the concentrations of IL7, IL10, IL12p70, and IL13 (Panel c). Instrumental variable analyses indicate that the inter-correlations between the cytokines match the causal effect estimates; this suggests a causal role of VEGF in mediating a cascade effect on IL7, IL10, IL12p70, and IL13 (Panel d).

To assess whether the genetic loci identified harbor additional independent variants, we performed conditional meta-analyses for each cytokine-lead SNP pairs. The 17q12 locus harbored five variants independently associated with macrophage inflammatory protein-1β (MIP1b) concentration **(Supplementary Table 5)**. These five statistically independent variants together explained 34% of the variation in circulating MIP1b. The variance explained by the independent variants for each cytokine are listed in **Supplementary Table 6**. The SNPs independently associated with VEGF explain 15% of the variation in circulating VEGF. The *MEP1B* locus had a large influence on circulating TRAIL and the variants identified here explain 14% of observed variance in TRAIL concentration. In addition, GROa was shown to be under a strong genetic control as the identified SNPs explained 10% of the circulating GROa variance.

### Expression analyses link three SNPs, cytokine mRNA and circulating cytokine concentrations

To further elucidate the molecular mechanisms underlying the genome-wide association peaks, we performed peripheral blood expression quantitative trait loci (eQTL) analyses for lead SNPs. Transcriptomic data were meta-analyzed for 2,177 participants from YFS and FINRISK2007. These analyses identified 26 significant eQTLs for 15 SNPs. The eQTLs for SNPs in **Table 1** are presented in **Table 2**; all significant eQTLs, including SNPs not listed in **Table 1**, are reported in **Supplementary Table 7**. Three SNPs were associated with both the mRNA encoding the cytokine and the pertinent circulating cytokine concentration. Allele rs4778636-A in *IL16* was associated with decreased concentration of circulating IL16 (**Table 1**), but increased concentration of *IL16* mRNA (**Fig. 3**). The Ensembl’s Variant Effect Predictor suggests that the A-allele creates a splice site variant leading to nonsense-mediated decay of *IL16* mRNA^18^. Increased *IL16* mRNA concentration may therefore be a compensatory mechanism to account for missing circulating IL16. The genome-wide association of rs4778636 with circulating IL16 confirms results from a small candidate gene study^19^. The MIP1b-associated variant (rs113877493) is located approximately 200kB away from a cluster of cytokine-coding genes in 17q21. According to our results, the MIP1b-associatied SNP is an eQTL for four of the genes (*CCL4L1*, *CCL4L2*, *CCL3L1*, *CCL3L3*) belonging to this cluster. The variant which was associated with concentration of PDGFbb and resided at a transcription factor binding site (rs13412535) was associated with expression of *SERPINE2*. Interestingly, rs13412535 is located within an intron of *SERPINE2* within the gene sequence instead of upstream from the gene, where transcription factors have been traditionally thought to bind.

**Figure 3.**
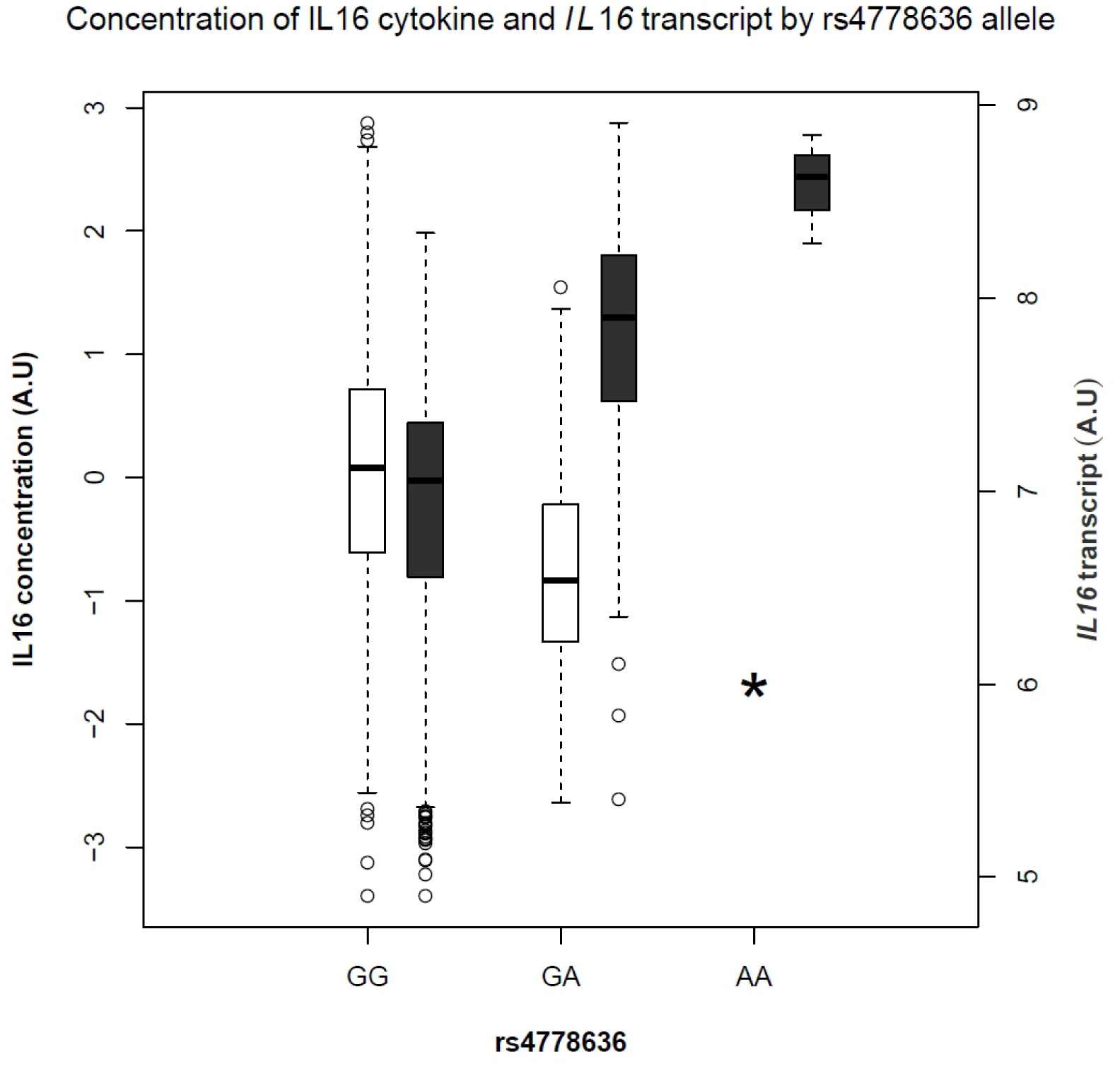
Boxplots of interleukin-16 (IL16) cytokine and *IL16* mRNA concentrations. The mRNA and circulating cytokine concentration are depicted by the dark grey and white boxes, respectively. Variant rs4778636 is located within *IL16* gene. The asterisk indicates undetectable concentration of circulating IL16. The A-allele of rs4778636 leads to non-sense mediated decay of *IL16* mRNA and thus circulating concentration of IL16 is undetectable in rs4778636-AA homozygotes.

**Table 2.**
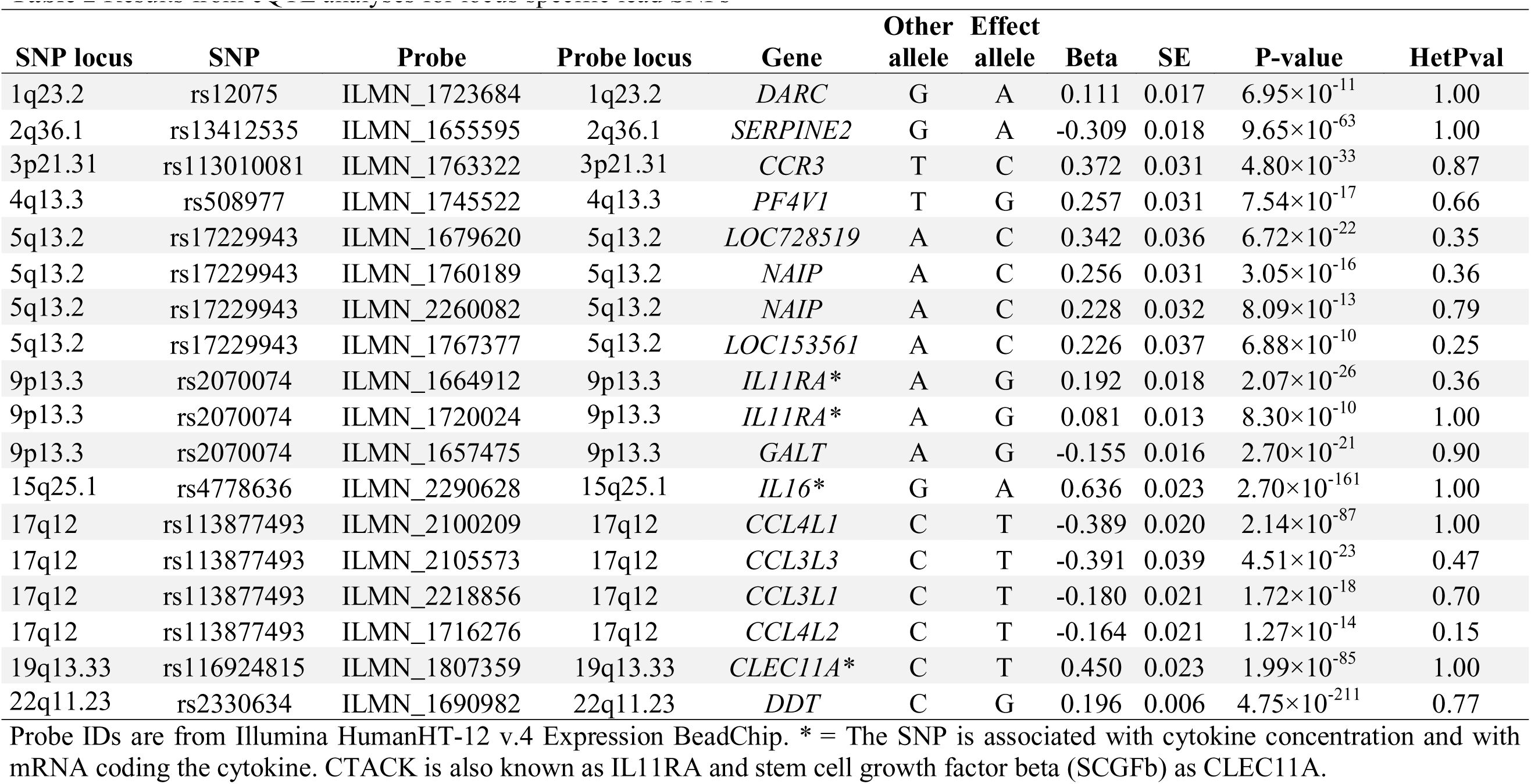
Results from eQTL analyses for locus specific lead SNPs

### Novel drug targets for celiac disease and Behcet’s disease and wider indication for IL2ra antagonists

To bridge the connection between the cytokine traits and disease, we searched the GWAS catalogue for SNP-trait and SNP-disease associations within a 1Mb window from cytokine lead SNPs^20^ (www.genome.gov/gwastudies). The search window approach was selected since many GWASes for the complex diseases have been conducted with HapMap2 and therefore missed variants present in the 1000 Genomes imputation panel^21^. The results for cytokine-linked SNPs previously associated with disease status are listed in **Table 3**; the corresponding associations for cytokine-linked SNPs previously associated with quantitative traits are listed in **Supplementary Table 8**. The circulating interleukin-2 receptor alpha subunit (IL2ra) increasing variant (rs12722489-C) has previously been shown to increase the risk for Crohn’s disease and multiple sclerosis^1,22^. The variant rs943072-G here associated with increased concentrations of VEGF and IL12p70 has previously been linked to increased risk for ulcerative colitis^23^. In addition, both cytokines have been shown to be increased in patients with an inflammatory bowel disease^24,25^. These results link the VEGF cascade depicted in Figure 2C to inflammatory bowel disease. Variant rs7616215-T here associated with increased concentration of MIP1b has previously been shown to decrease the risk for Behcet’s disease^26^. In addition to disease associations, the cytokine loci identified here were linked to 15 different molecular traits. For example, rs2251746 located at 1q23.2 within *FCER1A* and previously linked to IgE concentration was here associated with circulating concentration of GROa and MCP1.

**Table 3.**
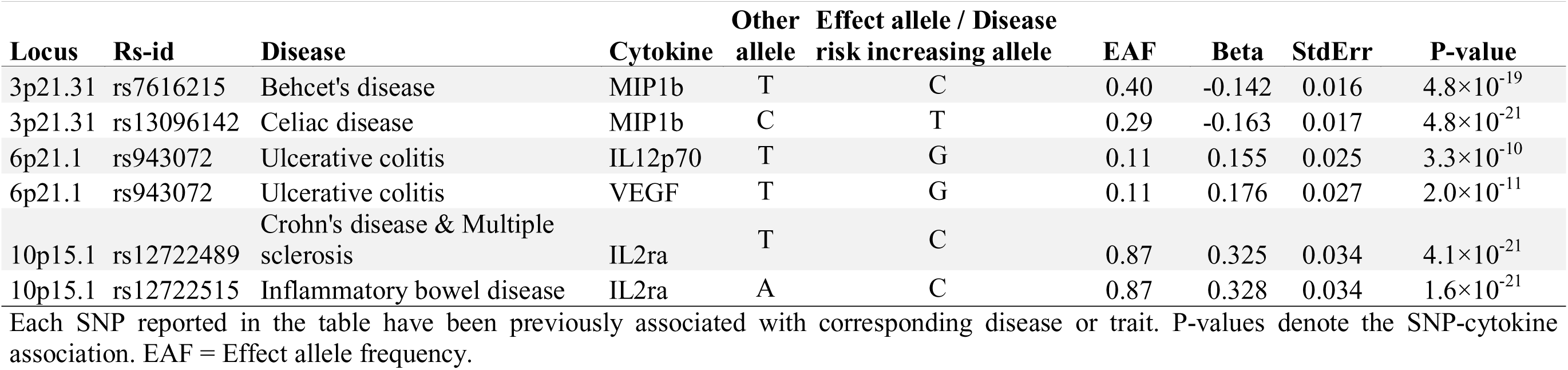
SNPs previously associated with disease and here found to be associated with cytokine concentrations.

## Discussion

This study identified 27 (17 novel) genetic loci contributing to circulating concentration of one or more cytokines in the general population. In three cases, the genomic analyses linked the genetic variant via mRNA to the circulating cytokine concentration, and therefore likely pinpointed the causal gene. Genetic evidence identify a potentially novel drug target for treatment of Behcet’s disease and celiac disease, since variants associated with decreased concentration of MIP1b in this study have previously been linked with increased risk of these diseases^26,27^. Furthermore, our results point to a novel indication for IL2ra-antagonist, since the IL2-ra variant identified in our study has previously been associated with increased risk of Crohn’s disease. The IL2ra-targeting antibody daclizumab reduces annualized relapse rate in multiple sclerosis patients by 50% compared to placebo^28^. Our results suggest that daclizumab might be beneficial for patients with Crohn’s disease as well.

The association of one locus to multiple circulating cytokines might help to understand clustering of autoimmune diseases. For example, a person with rheumatoid arthritis has more than 160-fold risk for autoimmune thyroiditis compared to the general population^5^. A genetic variant in *VEGFA* was here associated with increased circulating VEGF and shown to have a causal cascade effect on IL7, IL10, IL13 and IL12p70 (Fig 2C-D). Axitinib, a VEGF-receptor inhibitor, has been shown to reduce IL12p70 production in cultured monocyte-derived dendritic cells^29^, which support our findings. Furthermore, VEGFA-transcription follows the same temporal pattern with *IL7* transcription during wound healing, and it has been suggested that the secretion of these two cytokines might be connected by the same pathway^30^. These results from prior functional studies support our genetic evidence in humans that VEGF causally regulates secretion of IL7, IL10, IL13 and IL12p70 under normal physiological conditions in healthy individuals. VEGF contributes to pathogenesis of asthma by stimulating angiogenesis, edema, inflammation and airway remodeling. These effects are partly mediated via VEGF-mediated upregulation of IL13^31^. Most importantly, by linking the VEGF cascade with the risk for ulcerative colitis, our results suggest that drugs targeting VEGF might have potential in treatment of ulcerative colitis^23^. Furthermore, variants located 530 kB from rs10761731 (10q21.3; i.e. also associated VEGF like 6p21.1 ) were associated with ulcerative colitis^2^. The same region had suggestive association with age at diagnosis of inflammatory bowel disease in another immunochip-based study^32^. This is further substantiated by GWAS of inflammatory bowel diseases, where the identified loci were enriched in pathways related VEGF cascade cytokines (IL12 and IL10)^2^.

Another result with direct implications for drug development was found for the TRAIL-associated *MEP1B* locus. Drugs targeting TRAIL-signaling include Conatumumab, which is currently in phase II trials for treatment of a variety of cancers (ClinicalTrials.gov Identifier: NCT01327612). The low-frequency variant here identified to have a major influence on circulating TRAIL levels could help to clarify the causality of TRAIL-signaling in the development of cancer and assess potential side-effects from TRAIL-lowering similarly as done for IL1α/β and other targets^33^.

Plasma IL16 levels have previously been associated to HIV progression^34^, but the locus has not been linked to HIV-related traits in GWAS^35^. This might be due to the small allele frequency in Europeans or possible recessive effects. However, our results offer a possibility to examine HIV progression in human knock-outs, since the frequency of rs4778636-AA homozygote is up to 25% in the Yoruba population of Nigeria^36^. Circulating IL16 has been suggested to serve as a biomarker for impaired kidney transplant function, emphysema and interferon-β treatment efficacy in multiple sclerosis^37–39^. If IL16 is used as a biomarker, the relatively high frequency of rs4778636-AA homozygotes in African and Asian ethnicities must be accounted for.

Although TNFa antibodies have been successfully used to treat various autoimmune diseases^6,7^, no associations with cytokine concentrations were detected within the *TNF* locus (encoding TNF-α) in the MHC class III region at 6p21.33. The MHC region displays a unique pattern of linkage disequilibrium: a subset of MHC haplotypes have moderate correlation with each other whereas variants within same haplotype block form a stronger correlation structure^40^. This distinct linkage disequilibrium pattern may create complex epistasis to MHC locus, which might impede the association studies in this region. Furthermore, the cytokines were measured in conditions without stimulation. Measuring the cytokines concentrations after antigen stimulus might reveal additional loci contributing to circulating cytokine concentrations.

Variable degree of heterogeneity was found for of the 15 genetic loci identified. For most of the heterogenic loci, another SNP with genome-wide association but no heterogeneity could be found (see Supplemental note p. 111-124). The reason for occasional heterogeneity is likely differences in stability and accuracy of the cytokine experimental assay. In the case of the association between rs12075 and MCP1, differences in blood sample processing may also contribute: heparin treatment have been shown to release significant amounts of MCP1 from DARC, and thus affect the association signal caused by altered receptor binding properties^41^.

In conclusion, we identified 17 new loci contributing to the genetic regulation of circulating concentrations of cytokines. Improved understanding of the genetic basis of these inflammatory markers will help to clarify the causal roles of cytokine signaling and upstream inflammation in immune-related and other chronic diseases. By linking rs4778636 to undetectable concentration of IL16, these results enable studies of HIV and other inflammatory diseases in IL16 knock-out humans. In addition, we identified a potential novel drug target for treatment of Behcet’s disease and celiac disease as well as indicated a possibility to expand indication of daclizumab from multiple sclerosis to Crohn’s disease. These results provide the basis for further studies on the molecular regulation of the immune system in health and disease.

## Methods

### Study populations

#### The Cardiovascular Risk in Young Finns Study

The Cardiovascular Risk in Young Finns Study (YFS) is a multicenter follow-up study with randomly chosen subjects from Finnish cities of Helsinki, Kuopio, Oulu, Tampere, and Turku and their rural surroundings. The study began in 1980 when 3596 children and young adults participated to first cross-sectional survey. The follow-up visits have been conducted in 1983, 1986, 1989, 2001, 2007 and 2011. The present cross-sectional study includes 2,019 unrelated individuals who participated to 2007 follow-up and who had both cytokine measurements and genotype data available. In addition, gene expression data from 1,664 participants of the 2011 follow-up were analyzed for the present study. All participants gave written informed consent and the study was approved by local ethics committees^42^.

#### FINRISK

FINRISK surveys are population-based cross-sectional studies conducted every five years to monitor the levels of chronic disease risk factors in Finland. Each survey includes 25-74-year-old randomly chosen subjects from five geographical areas of Finland. The present study analyses cytokine data from participants of the 1997 and 2002 surveys. The study visit includes a clinical examination and semi-fasting blood sampling. For eQTL analyses, a peripheral blood sample was drawn to quantify mRNA expression profiles from a subset of 513 FINRISK2007 participants living in the Helsinki area^43^.

#### Cytokine quantification

From YFS and FINRISK2002, total of 48 cytokines were measured by using Bio-Rad’s premixed Bio-Plex Pro Human Cytokine 27-plex Assay and 21-plex Assay, and Bio-Plex 200 reader with Bio-Plex 6.0 software^44^. The assays were performed according to manufacturer’s instructions, except, that the amount of beads, detection antibodies and streptavidin-phycoerythrin conjugate were used with 50% lower concentrations than recommended. Only measures within the cytokine-specific detection range were included in the analyses. Cytokines with >90% of values missing were excluded (7 out of 48).

In FINRISK1997, a total of 17 cytokines overlapped with those measured in FINRISK2002 and YFS and were thus included in the GWAS. Genome-wide and cytokine data were available from up to 4608 FINRISK1997 participants and up to 1705 FINRISK2002 participants. The cytokine quantification was performed from EDTA plasma in FINRISK1997, from heparin plasma in FINRISK2002 and from serum in YFS.

#### Statistical analyses

Subjects whose cytokine concentration were below or above of laboratory analysis detection limits were omitted from the analyses. Cytokine distributions were first normalized with inverse transformation. The transformed phenotypes were then adjusted for age, sex, body mass index and ten first genetic principal components by calculating residuals of linear regression model. Subsequently, another inverse transformation was performed for model residuals to ensure normally distributed phenotypes. All effects sizes are hereby in SD-scaled units.

Genome-wide association testing were performed using Snptest2 software version 2.5beta^45,46^. Imputation inaccuracy was assessed with missing data likelihood score test. Allele effects were estimated from additive model (-frequentist 1). To prevent centering and scaling of phenotypes, use_raw_phenotypes option was enabled.

Meta-analyses were performed using METAL software (version 2011-03-25)^47^. Prior to metaanalysis all markers with imputation info <0.7, model fit info <0.7 and minor allele count <10 were excluded. After filtering, 10.7 million markers were included in the meta-analysis. Each association test was weighted by sample size. Genomic control correction was used to account for population stratification and cryptic relatedness. Heterogeneity statistics based on Cochrane’s Q-test were calculated for all markers included in meta-analysis to estimate heterogeneity of effect sizes across the three cohorts. Statistical significance was set at *P*<1.2×10^-9^, i.e. dividing the established genome-wide statistical significance (*P*<5×10^-8^) by 41 (number of cytokines analyzed). Random effect meta-analyses were performed with *R* software. To assess whether cytokine-associated loci harbor multiple variants independently associated with cytokine concentration, the models were further adjusted with locus-specific lead SNP and the model was fitted against the concentrations of all included cytokines. All independent SNPs were used to calculate the total variance explained by the identified loci. The proportion of total variance explained by independent SNPs was calculated by following formula:

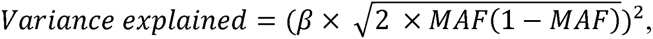

where β is the regression coefficient and MAF is minor allele frequency^48^.

Instrumental variable analysis of the VEGF cascade effects were performed with two-stage least squares regression using *ivreg* from the R-package AER^17^. A weighted gene score based on the sum of cytokine-increasing alleles of the two independent variants associated with VEGF (rs6921438 and rs12214617) was used as the instrument. The estimates from the three individual cohorts were combined with inverse variance weighted meta-analyses. The observational associations were calculated as linear regression between the cytokines after adjusting for age, sex, body-mass index, and the ten first genetic principal components.

#### eQTL analyses

Gene expression data from peripheral blood leukocytes were available from 1,664 participants from the YFS 30-year follow-up survey in 2011. After preprocessing the expression levels were analyzed with an Illumina HumanHT-12 version 4 Expression BeadChip. Raw Illumina probe data was exported from Beadstudio and analysed in *R* using the *Bioconductor* packages. The HT-12 v4 BeadChip contains 47,231 expression probes. Probes expressed by <5% of the participants were excluded, resulting in 19,637 probes^49^.

Gene expression data from peripheral blood leukocytes were analyzed for 518 individuals from FINRISK2007 with Illumina HumanHT-12 version 3 Expression BeadChips. Probes which were not autosomal, were complementary to cDNA from erythrocyte globin components and mapped to more than one genomic position were excluded. The filtering resulted to total of 35,419 probes included in the analyses^43^.

Linear regression with probe as a dependent variable was used to test associations between cytokine-associated SNPs and transcripts. Age and sex were used as covariates. Genotype dosage was calculated for each included SNP with Qctool software^50^. Probes with P-value <0.05 in both YFS and FINRISK07 were included in the eQTL meta-analyses. The eQTL results were combined by inverse variance weighted meta-analysis.

#### Database searches

To assess whether cytokine concentration could contribute to disease pathogenesis, we searched the GWAS catalogue for complex trait-associated SNPs within 1Mb window from the cytokine lead SNPs^20^. The catalogue was downloaded on July 6, 2015 and catalogue SNPs with *P*<5×10^-8^ were selected for further analyses. The statistical significance for SNP-cytokine association was inferred at *P* < 1.2×10^-9^.

## Acknowledgements

The Young Finns Study has been financially supported by the Academy of Finland: grants 286284 (T.L.), 134309 (Eye), 126925, 121584, 124282, 129378 (Salve), 117787 (Gendi), and 41071 (Skidi); the Social Insurance Institution of Finland; Kuopio, Tampere and Turku University Hospital Medical Funds (grant X51001 for T.L.); Juho Vainio Foundation; Paavo Nurmi Foundation; Finnish Foundation of Cardiovascular Research; Finnish Cultural Foundation; Tampere Tuberculosis Foundation (T.L.); Emil Aaltonen Foundation; and Yrjö Jahnsson Foundation. PW was supported by Finnish Diabetes Research Foundation, Novo Nordisk Foundation and Yrjö Jahnsson Foundation. VS was supported by the Academy of Finland, grant number 139635 and the Finnish Foundation for Cardiovascular Research.

We gratefully acknowledge Ville Aalto and Irina Lisinen for the expert technical assistance in the data management.

## Author contributions

AVA-O, PW, JK and OTR wrote the manuscript. KA, SJ, MM and MS performed the cytokine measurements. ER and IS acquired the expression data from YFS subjects and performed the quality control. APS performed the genotype imputation in all involved studies. ASH, NP, TL, MK, LPL, SR, AP, MP, JSV, VS gave critical comments regarding the manuscript. JK, OTR, TL, SJ and VS supervised the research. OTR and VS organized the data collections.

---

**Supplementary Figure 1** Pearson correlations between the circulating concentrations of different cytokines. The correlations are based on inverse transformed values and are adjusted for age, sex, ten first genetic principal components, and body mass index.

